# Chronically elevated FGF23 drives sustained renal ERK signaling and inflammatory transcriptional programs mitigated by cFGF23 gene therapy

**DOI:** 10.1101/2024.01.18.576116

**Authors:** Adrian Salas-Bastos, Claire Bardet, Klaudia Kopper, Louisa Jauze, Fanny Collaud, Amandine Francois, Matthias B. Moor, Gaozhi Chen, Moosa Mohammadi, Christian Stockmann, Lukas Sommer, Johannes Loffing, Giuseppe Ronzitti, Ganesh Pathare

**Affiliations:** Institute of Anatomy, University of Zurich, Zurich, Switzerland; Université de Paris, Institut des maladies musculo-squelettiques, Laboratory Orofacial Pathologies, Imaging and Biotherapies URP2496 and FHU-DDS-Net, Dental School, and Plateforme d’Imagerie du Vivant (PIV), Montrouge, France; Swiss National Centre of Competence in Research “Kidney Control of Homeostasis”, Switzerland; Genethon, 91000 Evry, France; Université Paris-Saclay, Univ Evry, Inserm, Genethon, Integrare research unit UMR_S951, 91000 Evry, France; Department of Nephrology and Hypertension, Inselspital University Hospital, Bern, Switzerland; Oujiang Laboratory (Zhejiang Lab for Regenerative Medicine, Vision, and Brain Health), School of Pharmaceutical Sciences, Wenzhou Medical University, Wenzhou, China; Institute of Cell Growth Factor, Oujiang Laboratory (Zhejiang Lab for Regenerative Medicine, Vision, and Brain Health), Wenzhou, China; Bone-Kidney Axis and Regeneration Laboratory, Department of Infectious Diseases and Public Health, Jockey Club College of Veterinary Medicine and Life Sciences, City University of Hong Kong, Hong Kong SAR

**Keywords:** FGF23, Klotho, X-linked hypophosphatemia (XLH), ERK1/2, inflammatory transcriptome

## Abstract

Fibroblast growth factor 23 (FGF23) levels are highly elevated in patients with chronic kidney disease (CKD); however, whether it serves merely as a biomarker or actively contributes to kidney inflammation remains unclear. Full-length FGF23 is cleaved into C-terminal FGF23 (cFGF23), which acts as a natural antagonist of FGF23 by inhibiting its binding to the FGF receptor (FGFR) and the co-receptor Klotho. Here, we show that chronically elevated FGF23 levels in a mouse model Hyp-Duk cause sustained-ERK signaling and an inflammatory and immune responses in the kidney. cFGF23, delivered via adeno-associated virus (AAV) gene therapy, successfully mitigated renal sustained-ERK signaling and inflammatory and immune responses in Hyp-Duk mice. On the other hand, acute physiological FGF23 levels *in vitro* elicited transient-ERK signaling with the expression of canonical early-ERK targets (*EGR1, JUNB, FOSB*). Consistent with *in vivo* findings, prolonged pathological FGF23 treatment *in-vitro* revealed sustained-ERK signaling with upregulation of late-ERK targets (*ETV4/5, SPRED1/2, SPRY2/4*) and unique inflammatory and immune gene signatures. These effects were significantly mitigated by FGFR and ERK inhibitors, as well as by recombinant cFGF23 treatment. In summary, chronically high levels of FGF23 induce kidney inflammation through the FGFR-Klotho complex and sustained-ERK activation, which are successfully mitigated by cFGF23 gene therapy.

**TRANSLATIONAL STATEMENT:** Chronically elevated circulating levels of Fibroblast Growth Factor 23 (FGF23) are strongly associated with inflammation and adverse outcomes in chronic kidney disease, but the underlying mechanisms remain poorly understood. Our study identifies sustained activation of the ERK signaling pathway as a key mechanism by which chronic FGF23 elevation induces inflammatory transcriptional programs in the kidney. Importantly, gene therapy using the C-terminal fragment of FGF23 (cFGF23) mitigates these effects, highlighting a potential therapeutic strategy to counteract the excessive FGF23 signaling.

## INTRODUCTION

The phosphaturic hormone fibroblast growth factor 23 (FGF23) is primarily produced by osteoblasts and osteocytes^1–3^. FGF23 mediates its phosphaturic effects by inducing asymmetric dimerization of its cognate FGF receptors (FGFR) in the kidneys in a Klotho and heparan sulfate (HS) dual coreceptor fashion^2,4–6^. The 1:2:1:1 FGF23-FGFR-Klotho-HS signaling complex rapidly triggers ERK signaling thus inducing expression of immediate early genes such as *Egr1, Fos* and *Jun,* necessary for regulating mineral metabolism^6–8^. Klotho is a single pass transmembrane protein and is primarily expressed in the kidneys as mice with a kidney-specific Klotho ablation have a similar phenotype as global Klotho knockout mice^9,10^. Klotho is also expressed in in the brain, parathyroid glands, and gonads, albeit in lower levels compared to the kidney^11^. Extracellular shedding results in soluble form of Klotho (sKlotho) that is found in blood, urine, and cerebrospinal fluid^10,12^. sKlotho in supraphysiologic levels can mimic some of the effects of transmembrane Klotho^6,13^. FGF23’s bioactivity is principally regulated by a proteolytic cleavage, resulting in inactive larger N-terminal (nFGF23) and C-terminal fragments (cFGF23). Intact (iFGF23 or simply FGF23), nFGF23 and cFGF23 coexist in the circulation^14,15^.The isolated cFGF23 peptide inhibits FGF23-Klotho binding thereby suppressing FGF23-mediated regulation of mineral metabolism^16,17^.

FGF23 levels are massively elevated in chronic kidney disease (CKD) patients and have been associated with inflammation, cardiovascular disease, morbidity and mortality^18–22^. The fact that chronic administration of cFGF23 alleviated renal and cardiovascular pathobiology in a rodent model of CKD lends credence to the notion that the association between high FGF23 and adverse events is more than a mere association but in fact signifies causality^23^. Whether elevated FGF23 causes kidney inflammation in CKD is not clear^24^. Inflammatory cytokines such as IL-6 and TNF-α are often elevated in CKD patients and may upregulate FGF23 expression in bone, as well as in immune cells and tissues that do not normally express FGF23^18,25–27^. Conversely, elevated FGF23 has been shown to induce inflammation in hepatocytes and immune cells, which do not express Klotho^26,28–30^. The Klotho-independent actions of FGF23 contradict the functional and structural data that proposes the essential role of Klotho in FGF23 function^5,6,13^.

FGF23 is elevated in numerous human genetic diseases of hypophosphatemia, including X-linked hypophosphatemia (XLH), which is caused by inactivating mutations in the *PHEX* gene^31,32^. The clinical symptoms of XLH are growth retardation, leg bowing, rickets, osteomalacia, bone pain, and dental illness. Many of these symptoms are phenocopied in various murine models of XLH, whether arising from a spontaneous mutation such as the *Hyp-Duk* mouse or generated by inactivating the *Phex* gene^17,31,33^. To date, no inflammatory lesions have been reported in the kidneys or heart of XLH patients and mouse models who have lifelong elevated FGF23 levels^33–36^. However, recently, one study found that XLH patients have elevated circulatory inflammatory markers, suggesting a possible link between chronic FGF23 elevation and inflammation in XLH^37^.

Physiological regulators like dietary phosphate, vitamin-D and other hormones stimulate FGF23 transiently within a narrow physiologic range^38^. In contrast, CKD and XLH patients have chronically highly elevated FGF23 levels^18,19,21,22,38^. The differential effects of low-transitory (∼physiological) versus high-prolonged (∼pathological) FGF23 levels on temporal phases of ERK activation are unclear. Continued FGF23 availability for FGFR-Klotho binding in the kidney may lead to sustained-ERK activity, but this has not been demonstrated yet. Sustained-ERK activation regulates several transcription factors and gene signatures that govern inflammation and immune responses^39,40^. We hypothesize that chronically elevated FGF23 levels may cause inflammation and immune gene signatures via sustained-ERK signaling, expanding from the canonical role of FGF23 as a regulator of mineral metabolism.

## RESULTS

### 1. Chronically elevated FGF23 levels in mice induce sustained-ERK signaling, inflammation and immune gene signatures in the kidneys, which are mitigated by cFGF23 gene therapy

An adeno-associated virus (AAV) vector was generated to secrete human cFGF23 from the liver and increase circulating cFGF23 (AAV-cFGF23)^17^. Fig. 1A illustrates the experimental approach used in this study. As expected, serum FGF23 levels in Hyp-Duk mice were significantly higher than in WT mice (Fig. 1B). Circulating human cFGF23 levels were significantly increased in AAV-cFGF23-treated Hyp-Duk mice (Fig. 1C), while this assay did not detect endogenous mouse cFGF23 due to species specificity (Fig. S1). RNA-seq was performed on kidney samples of WT, Hyp-Duk and Hyp-Duk mice treated with AAV-cFGF23 (cFGF23-Hyp-Duk).

**Fig. 1.**
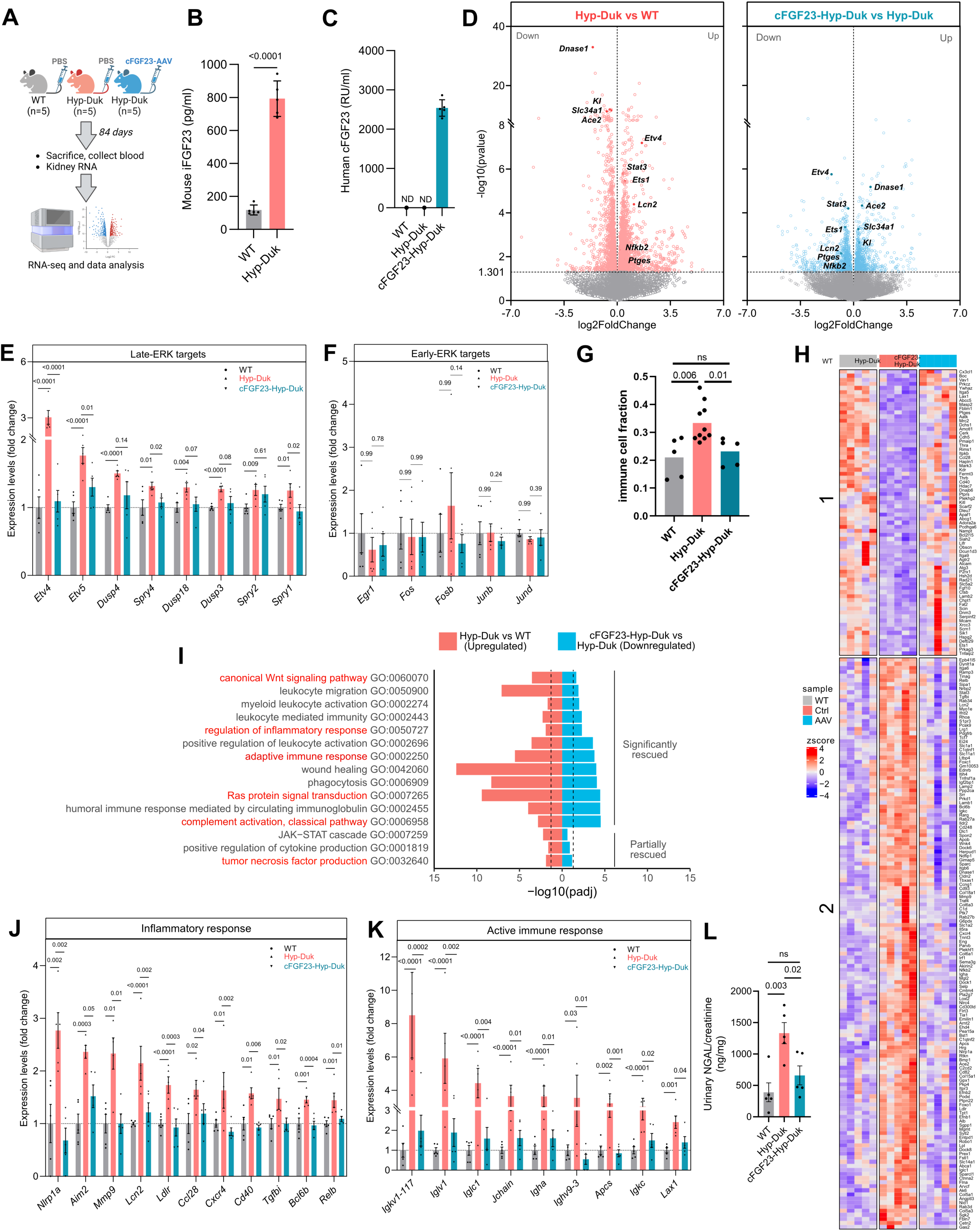
Elevated FGF23 levels in Hyp-Duk mice induces sustained-ERK signaling and inflammatory, and immune gene signatures in the kidneys, which are mitigated by cFGF23 gene therapy. **A)** Graphical illustration showing the experimental design for the *in vivo* experiment. **B)** Mouse iFGF23 levels in the serum of WT and Hyp-Duk mice (n=5, each group). **C)** Human cFGF23 levels in the serum of WT, Hyp-Duk, and Hyp-Duk mice receiving AAV-FGF23 (cFGF23-Hyp-Duk); n=5, each group; ND: not detected. **D)** Volcano plots illustrating DEGs identified by RNA-seq in the kidneys of WT, Hyp-Duk, and cFGF23-Hyp-Duk mice. *Left panel*: DEGs compared between Hyp-Duk versus WT; *right panel*: DEGs compared between cFGF23-Hyp-Duk versus Hyp-Duk. Note that numerous DEGs identified in the kidneys of Hyp-Duk were significantly oppositely regulated in the kidneys of cFGF23-Hyp-Duk mice. **E&F)** Gene expression levels of late- and early-ERK targets in the kidneys of WT, Hyp-Duk, and cFGF23-Hyp-Duk mice. **G)** Fraction of transcriptome-inferred immune-like cells in kidneys of WT, Hyp-Duk, and cFGF23-Hyp-Duk mice, as estimated by bulk RNA-seq deconvolution using reference kidney single-cell transcriptomes. **H)** Heatmap of ImmPort DEGs in the kidneys of Hyp-Duk rescued by AAV-cFGF23 treatment (block 1 and 2 represent downregulated and upregulated genes in Hyp-Duk mice, respectively. **I)** Selected significantly upregulated GO pathways (biological processes) in the kidneys of Hyp-Duk versus WT mice. Note that these pathways were either significantly or partially rescued in the kidneys of cFGF23-Hyp-Duk mice. Dotted line indicates threshold for *padj<0.05*. The pathway names in red indicate specific signal transduction pathways implicated in inflammatory and immune gene signatures. **J&K)** Graphs illustrating representative genes implicated in GO pathways of overall inflammatory and immune responses in the kidneys of WT, Hyp-Duk, and cFGF23-Hyp-Duk mice. **L)** Urinary NGAL levels normalized to creatinine. n=5, each group. All values are expressed as arithmetic means ± SE.

The number of DEGs (p-value<0.05) in the kidneys of Hyp-Duk and cFGF23-Hyp-Duk are shown in Fig. S2 (tables S1/2). Volcano plots illustrate DEGs between the kidneys of Hyp-Duk versus WT and AAV-cFGF23-Hyp-Duk versus Hyp-Duk mice (Fig. 1D). Notably, many canonical FGF23 targets, such as *Dnase1*, *Slc34a1*, *Kl*, and *Ace2*, were downregulated in the kidney of Hyp-Duk mice but upregulated in the kidneys of cFGF23-Hyp-Duk mice, validating the gene therapy approach (Fig. 1D). Previously unrecognized ERK and inflammatory targets, such as *Etv4*, *Stat3*, *Ets1*, and *Nfkb2*, were significantly upregulated in the kidneys of Hyp-Duk mice but downregulated the kidneys of cFGF23-Hyp-Duk mice (Fig. 1D). Notably, only late-ERK targets were significantly upregulated in the kidneys of Hyp-Duk mice and completely or partially rescued in the kidneys of cFGF23-Hyp-Duk mice (Fig. 1E/S3). Early-ERK targets remained unchanged in the kidneys of Hyp-Duk mice (Fig. 1F).

To determine whether sustained FGF23 signaling alters the renal immune cell landscape, we performed bulk deconvolution of our renal RNA-seq data using reference kidney single-cell transcriptomes. The fraction of transcriptome-inferred immune-like cells was increased in the kidneys of Hyp-Duk mice compared with controls (Fig. 1G). In contrast, cFGF23 overexpression significantly reduced the immune-like cell fraction (Fig. 1G). Further, we intersected DEGs from Hyp-Duk kidneys with the immunological database ImmPort. We found that 20.5% of all DEGs corresponded to ImmPort-annotated genes (Fig. S4; left panel). Notably, cFGF23 gene therapy rescued 27.4% of these affected DEGs (Fig. S4; right panel), which are highlighted in the heatmap (Fig. 1H). GO enrichment analysis identified 277 pathways significantly upregulated (padj < 0.05) in Hyp-Duk kidneys and rescued in cFGF23-Hyp-Duk kidneys (tables S3/4/5). Many of these pathways were related to inflammation and immune responses. For example, ERK-mediated *Ras signaling*, *inflammatory and immune responses*, *Wnt signaling*, and *complement activation* were significantly upregulated in Hyp-Duk kidneys and rescued by cFGF23 gene therapy (Fig. 1I; tables S3/4/5). In contrast, the inflammatory *JAK-STAT cascade* and *TNF cytokine production* were significantly upregulated and only partially rescued (Fig. 1I; tables S3/4/5). This list is not exhaustive; the full table is provided in tables S3/4/5.

Fig. 1J/K show selected genes implicated in inflammatory and immune gene signatures that were upregulated in the kidneys of Hyp-Duk mice and reversed in the kidneys of cFGF23-Hyp-Duk mice. Genes involved in paracrine FGF signaling, such as *Fgf10*, *Hspg2*, and *Fgf2*, were significantly upregulated in the kidneys of Hyp-Duk mice and rescued in the kidneys of cFGF23-Hyp-Duk mice (Fig. S5). We found canonical mineral metabolism targets of FGF23 in the kidneys of Hyp-Duk mice, such as *Cyp27b1*, *Cyp24a1*, *Slc34a1*, and *Kl*, as well as several previously unrecognized gene targets, including *Slc34a2*, *Pth1r*, and *Calb1* (Fig. S6). Fig. S7 summarizes the pathways upregulated in the renal transcriptional signature associated with skeletal defects in Hyp-Duk mice. *Lcn2* upregulation in Hyp-Duk kidneys prompted us to measure urinary neutrophil gelatinase-associated lipocalin (NGAL), a marker of kidney inflammation and injury. Urinary NGAL levels were significantly elevated in Hyp-Duk mice and were rescued following AAV-cFGF23 treatment (Fig. 1L).

### 2. Sustained-ERK activation in vitro is induced by prolonged high, but not acute low, FGF23 concentrations

HEK293 (HEK^Wt^) cells endogenously express FGFR1-4 but not Klotho. We stably transfected these cells with Klotho to generate a stable Klotho HEK293 cell line (HEK^Kl^). HEK^Wt^ and HEK^Kl^ cells are excellent models for studying Klotho-independent and Klotho-dependent FGF23 functions, respectively^13,41^. The activation of pERK1/2 by FGF23 in the presence of Klotho was confirmed (Fig. S9). Treatment with 0.5 nM FGF23, which resembles the physiological condition, in HEK^Kl^ cells, resulted in a transient increase in pERK1/2 that was visible after 15 minutes but not at 24 hours (Fig. 2A, left panel). Treatment with 10 nM FGF23, a higher concentration mimicking the pathological levels, led to pERK1/2 activation at 15 minutes which persists to 24 hours (Fig. 2A, right panel). FGF23 concentrations exceeding 5 nM resulted in sustained pERK1/2 activation (Fig. 2B). To determine the precise time points for the transcriptomics analysis, we used both low and high FGF23 concentrations and measured *EGR1* mRNA as a canonical marker for FGF23 signaling. Upon 0.5 nM FGF23 treatment in HEK^Kl^ cells, *EGR1* mRNA peaked at 1 hour, with no statistically significant difference at 24 hours compared to the control HEK^Wt^-treated cells (Fig. 2C). Notably, 10 nM FGF23 treatment followed almost a similar trend except a 13-fold higher *EGR1* levels were still observed after 24 hours (Fig. 2D/E). Both pERK1/2 and *EGR1* levels remained unchanged in HEK^Wt^ cells throughout the treatment periods indicating the absolute requirement of the co-receptor Klotho for FGF23 signaling.

**Fig. 2.**
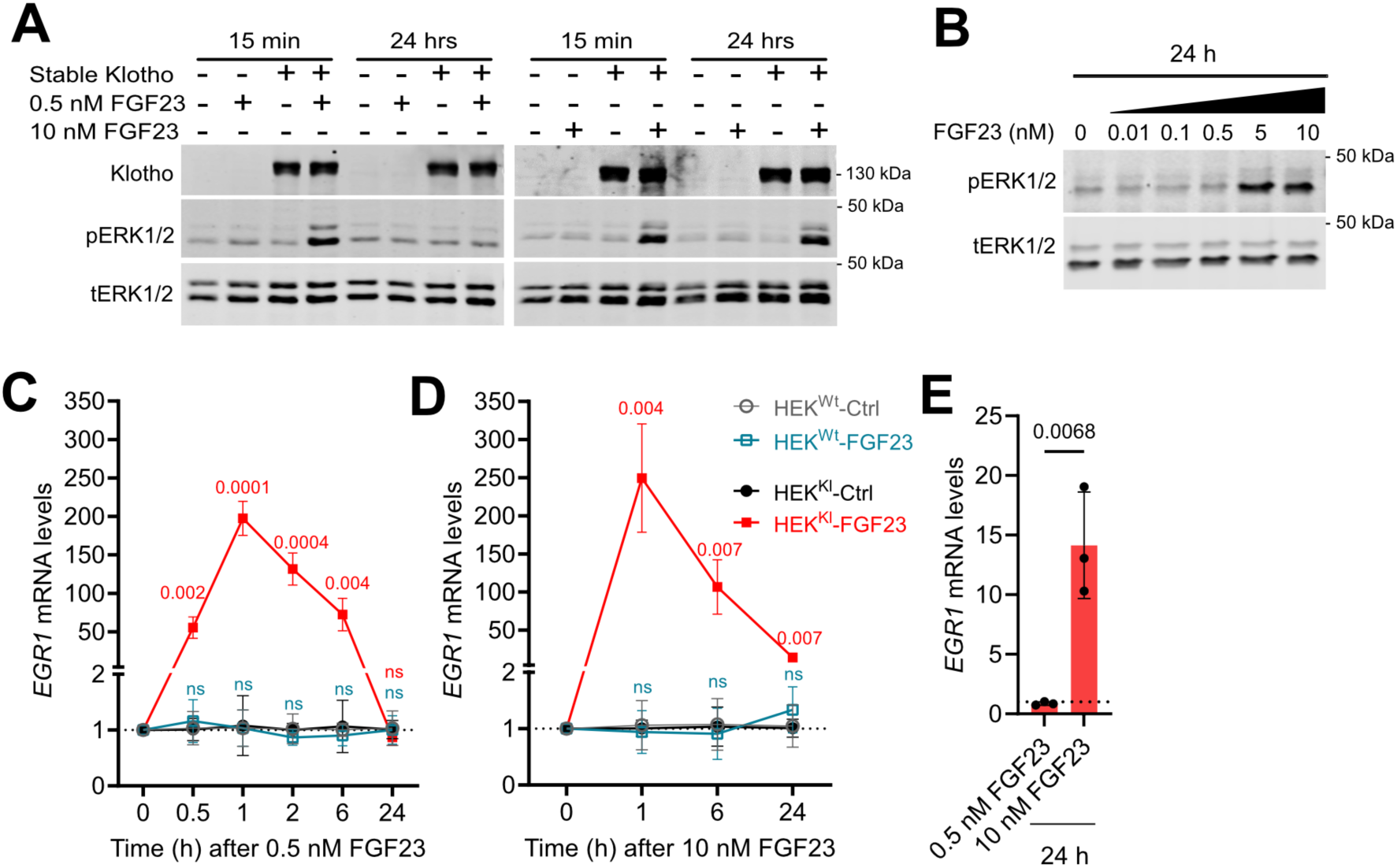
Effects of low versus high FGF23 concentrations (for 15 mins vs. 24 h) on ERK activation and *EGR1* levels. **A)** Original immunoblots of Klotho, pERK1/2 and tERK1/2 upon treatment with 0.5 nM and 10 nM FGF23 in HEK^Kl^ and HEK^Wt^ cells for 15 min and 24 hrs. **B)** Original immunoblots of pERK1/2 and ERK in HEK^Kl^ cells upon treating with increasing concentrations of FGF23 for 24 hours. **C&D)** Fold change *EGR1* mRNA levels, normalized to *GAPDH* upon treating 0.5 nM and 10 nM FGF23 in HEK^Kl^/HEK^Wt^ cells over 24 hours, respectively (n=3). *p* values in red and teal indicate comparisons between FGF23 vs control treatment in HEK^Kl^ and HEK^Wt^ cells, respectively. ns=*p*>0.05. **E)** *EGR1* mRNA, normalized to *GAPDH,* upon treating with 0.5 nM and 10 nM FGF23 for 24 hours in HEK^Kl^ cells. n=3, each group. All the values are expressed as arithmetic means ± SE.

### 3. Chronically elevated FGF23 levels induce sustained-ERK signaling and inflammation-related gene signatures *in vitro*

To evaluate the transcriptomic profile after treatment with physiological or pathological FGF23 concentrations, we performed RNA-seq analysis as illustrated in Fig. 3A. Physiological FGF23 treatment in HEK^Wt^ cells did not affect the transcriptome (Fig. 3B, left panel; table S6; and Fig. S10A). Physiological FGF23 treatment in HEK^Kl^ cells led to 64 differentially expressed genes (DEGs), with 61 upregulated and 3 downregulated (Fig. 3B, right panel; table S7). Canonical early genes such as *EGR1, FOS* and *JUNB* were among the top upregulated genes, validating our approach. Although pathological FGF23 concentrations in HEK^Wt^ cells resulted in few DEGs, these showed log2FC<1, suggesting a minimal non-Klotho-dependent effect of FGF23 (Fig. 3C, left panel; table S8). While pathological FGF23 treatment in HEK^Kl^ cells demonstrated a robust transcriptomics response (Fig. 3C, right panel; table S9).

**Fig. 3.**
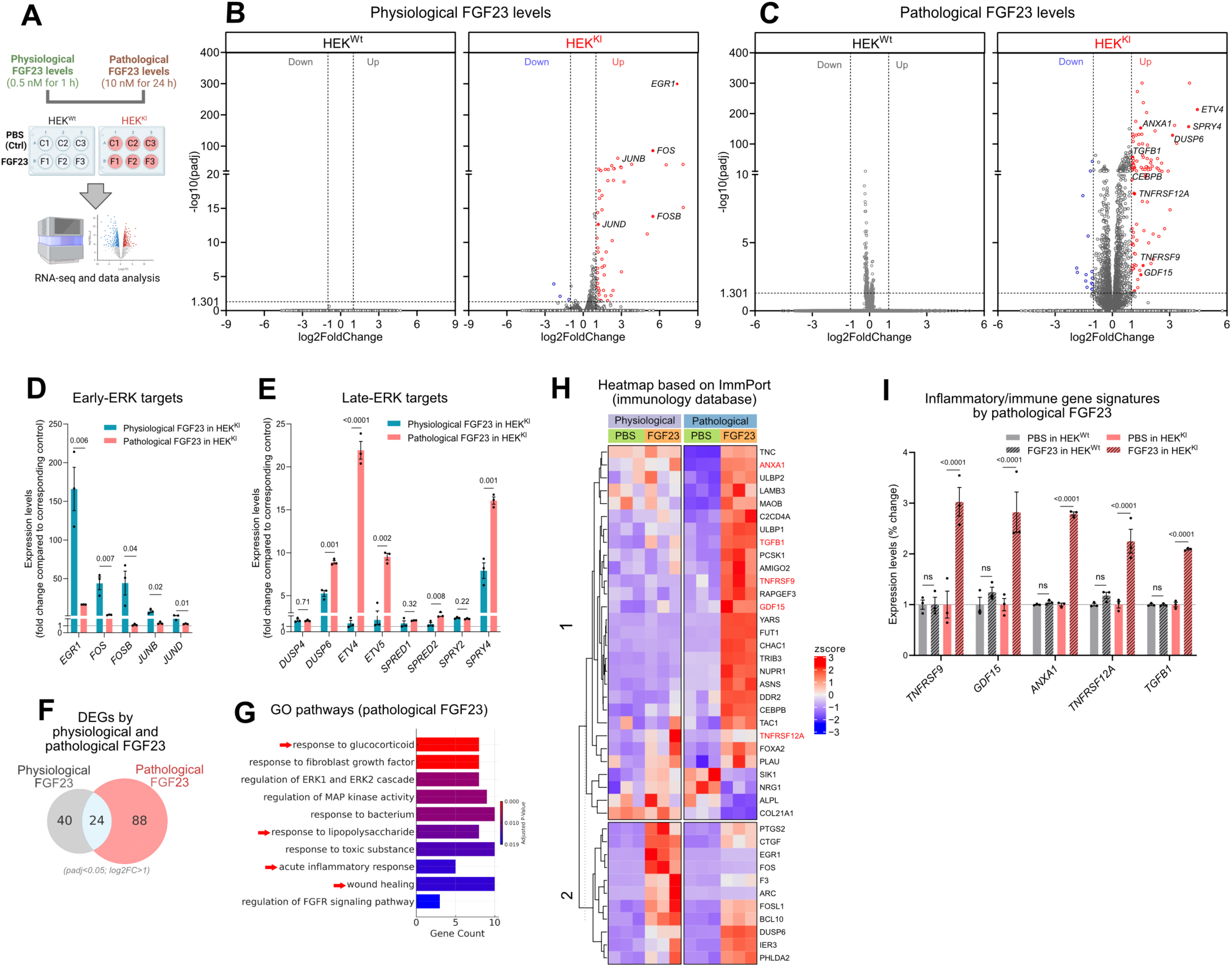
Identification of temporal ERK signaling by physiological versus pathological FGF23 concentrations. **A)** Graphic illustration showing the experimental design for RNA-seq. **B&C)** Volcano plots illustrating all the genes identified by RNA-seq upon physiological and FGF23 pathological in HEK^Wt^/HEK^Kl^ cells, respectively. The upregulated and downregulated genes (abs(log2FC)>1) are shown in red and blue, respectively. **D)** ERK targets that are statistically significantly upregulated by physiological FGF23 compared to pathological FGF23 in HEK^Kl^ cells. The gene expression is compared to the corresponding control. **E)** ERK targets that are either statistically significantly unchanged or upregulated by pathological FGF23 compared to physiological FGF23 in HEK^Kl^ cells. The gene expression is compared to the corresponding control. **F)** Venn diagram showing DEGs compared between physiological FGF23 and pathological FGF23 in HEK^Kl^ cells (padj<0.05; (abs(log2FC)>1). **G)** Selected upregulated GO pathways in biological processes enriched upon pathological FGF23 in HEK^Kl^ cells (padj<0.05; log2Fc>1). The red arrows indicate pathways implicated in inflammatory and immune gene signatures. **H)** Heatmap highlighting DEGs by pathological FGF23 in HEK^Kl^ cells that were present in the ImmPort database. Block 1 represents DEGs only affected by pathological FGF23, while block 2 represents DEGs affected by both pathological and physiological FGF23. Exemplary immune/inflammatory genes reported in literature are shown in red. **I)** Representative transcripts implicated in inflammatory and immune gene signature and upregulated only by pathological FGF23 in HEK^Kl^ cells (red bars). Note that these genes are unchanged by similar treatment in HEK^Wt^ cells (grey bars). (n=3), ns=padj>0.05. All the values are expressed as arithmetic means ± SE.

The physiological and pathological FGF23 treatments in HEK^Kl^ cells upregulated several ERK target genes (Fig. S11). Early immediate genes like *EGR1, FOS, FOSB, JUNB* and *JUND* were significantly upregulated by physiological FGF23 compared to pathological FGF23 treatment *(early-ERK targets;* Fig. 3D and S11). *EGR1* expression was 150-fold higher after physiological FGF23 treatment, but only 10-fold higher in pathological conditions, indicating its primary role in transient-ERK activity. In contrast, *JUNB* and *FOSB* were unchanged by pathological FGF23 treatment, suggesting they are strictly *early-ERK targets* (Fig. S11). Conversely, ERK targets such as *DUSP4/6*, *ETV1/4/5*, *SPRED1/2*, and *SPRY2/4* were either unchanged or significantly upregulated by pathological FGF23 compared to physiological FGF23 treatment (Fig. 3E, S12). These transcripts were identified as *late-ERK targets*. Among these, *ETV1/4/5* and *SPRED1/2* were unchanged by physiological FGF23, indicating they are strictly *late-ERK targets* (Fig. S12).

As depicted in the Venn diagram, 88 and 40 DEGs were exclusively regulated by pathological and physiological FGF23, respectively, with 24 DEGs overlapping (Fig. 3F). This suggests that physiological and pathological FGF23 levels drive divergent transcriptomic responses. Analysis of the Gene ontology (GO) pathways of biological processes linked to genes upregulated by pathological FGF23 identified several processes implicated in inflammation and immune response (Fig. 3G, red arrows. A complete list of GO pathways is available in table S11). Therefore, we intersected DEGs regulated by pathological FGF23 with the immunological database ImmPort (https://www.immport.org). This analysis retrieved a set of genes linked to inflammatory and immune signatures, which are highlighted in the heatmap (Fig. 3H). Block 1 of the heatmap shows the inflammatory and immune gene signatures exclusively regulated by pathological FGF23, while block 2 shows genes regulated by both pathological and physiological FGF23. Representative inflammatory and immune gene signatures (*TNFRSF9/12A*, *GDF15*, *ANXA1*, and *TGFB1*) by pathological FGF23 (block 1) are highlighted in Fig. 3I. Notably, the early- and late-ERK targets, as well as novel FGF23 targets implicated in inflammatory and immune gene signatures, are upregulated only in HEK^Kl^ cells and not in HEK^Wt^ cells (Fig. 3I; Fig. S11/S12).

### 4. HEK^Kl^ cells model renal transcriptional programs downstream of FGF23-Klotho signaling

To determine whether the transcriptomic responses observed *in vitro* recapitulate the renal gene expression changes in Hyp-Duk mice, we compared DEGs implicated between pathological FGF23-treated HEK^Kl^ cells and the kidneys of Hyp-Duk mice. Heatmap analysis of late-ERK targets revealed concordant upregulation of *ETV1*, *ETV4*, *ETV5*, *SPRED2*, *SPRY2*, and *SPRY4* in pathological FGF23-treated HEK^Kl^ cells and their murine orthologs in Hyp-Duk kidneys compared to their respective controls (Fig. 4A). Similarly, genes implicated in inflammatory and immune responses, including *NFKB1*, *RELB*, *STAT3*, *TNFRSF1A*, *CD44*, *MMP2*, *FLI1*, *ITGB1*, *ITGA5*, *ITGA6*, and *BCL2*, were concordantly upregulated in pathological FGF23-treated HEK^Kl^ cells and Hyp-Duk kidneys (Fig. 4B).

**Fig. 4.**
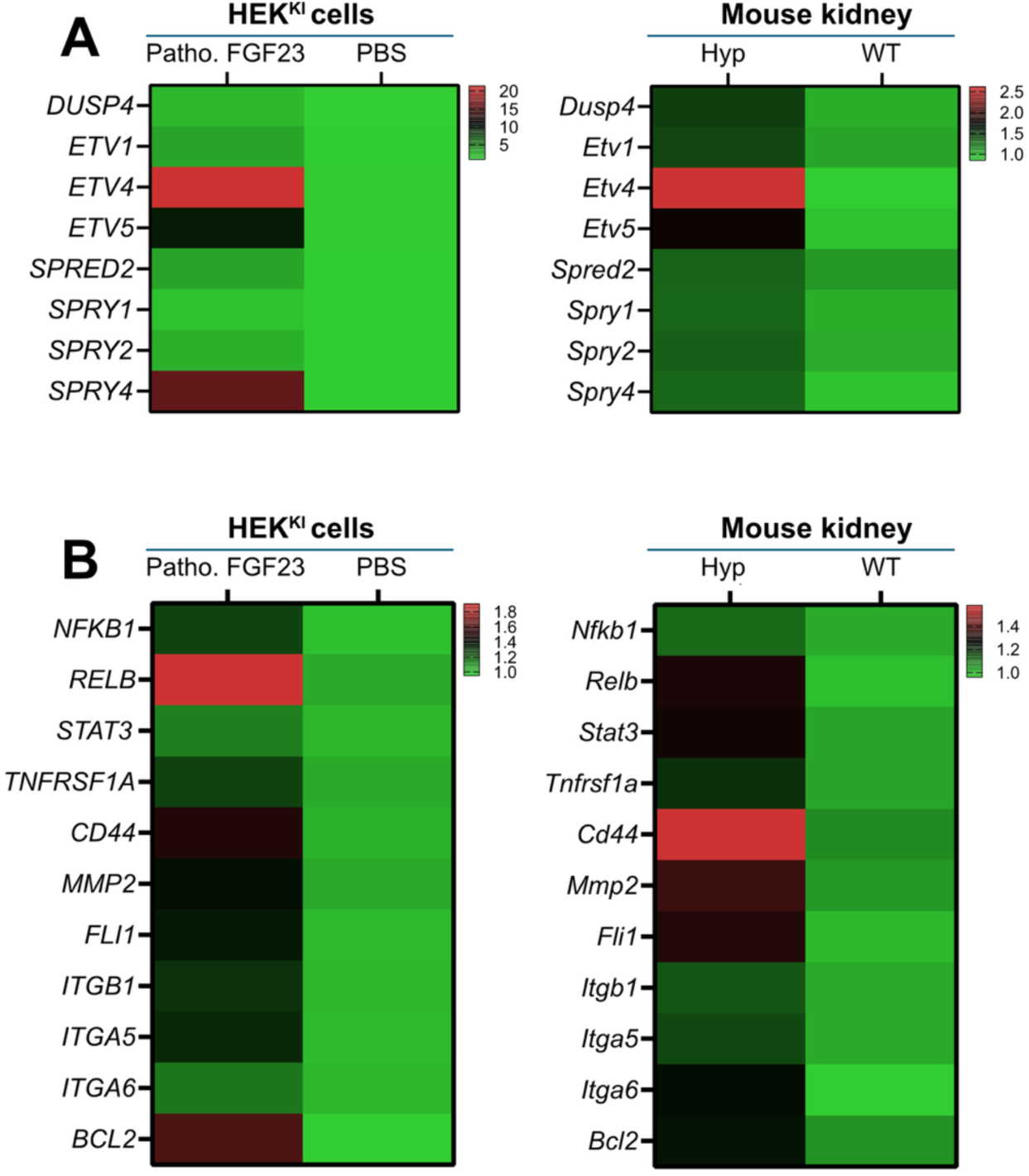
HEK^Kl^ cells model renal transcriptional programs downstream of FGF23-Klotho signaling. **(A)** Heatmaps showing relative expression of late ERK-responsive genes in HEK^Kl^ cells treated with pathological FGF23 compared with PBS control (n = 3 per group; P < 0.05) and in kidneys from Hyp-Duk (Hyp) versus wild-type (WT) mice (n = 5 per group; P < 0.05). **(B)** Heatmaps showing relative expression of inflammatory and immune-related genes in HEK^Kl^ cells (Patho. FGF23 vs. PBS; n = 3 per group; P < 0.05) and Hyp-Duk versus WT kidneys (n = 5 per group; P < 0.05).

### 5. FGF23-mediated inflammatory and immune gene signatures are mediated by FGFR and ERK in vitro

We used blockers of FGFR (PD173074) and MEK (U0126) to investigate the roles of FGFR and ERK in the effects mediated by pathological FGF23. The mRNA levels of newly identified targets were quantified using qRT-PCR. Pretreatment with either FGFR or MEK inhibitors completely reversed the FGF23-mediated upregulation of the early-ERK target *EGR1* (Fig. 5A, left panel). The late-ERK target *ETV4* was also entirely blocked by the FGFR inhibitor, while it was partially but significantly blocked by the MEK inhibitor (Fig. 5A, right panel). Next, we quantified the inflammation and immune targets such as *TNFRSF9/12A, GDF15, ANXA1,* and *TGFB1*. FGF23-mediated upregulation of mRNA levels of these genes were completely abrogated by FGFR inhibitors and partially but significantly reversed by MEK inhibitors (Fig. 5B). This implies that there is divergence of signaling post FGFR to different pathways.

**Fig. 5.**
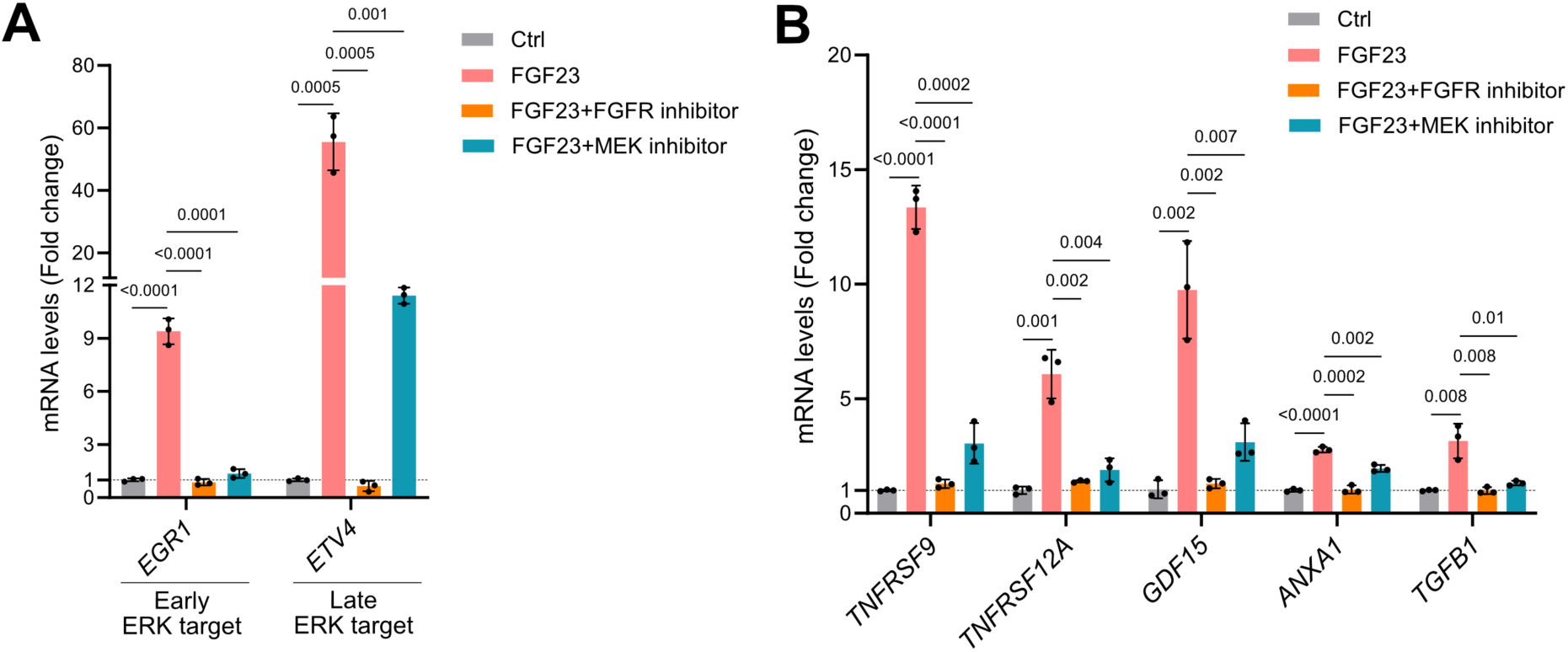
Pathological FGF23-mediated ERK signaling, inflammatory and immune gene signatures are inhibited by FGFR and MEK blockers. **A)** *EGR1* and *ETV4* mRNA, normalized to *GAPDH*, upon treating HEK^Kl^ cells with 10 nM FGF23 for 24 hours with or without FGFR inhibitor (PD173074) or MEK inhibitor (U0126) (n=3). **B)** mRNA levels of *TNFRSF9*, *TNFRSF12A*, *GDF15*, *ANXA1*, and *TGFB1* normalized to *GAPDH*, upon treating HEK^Kl^ cells with pathological FGF23 (10 nM FGF23 for 24 hours) with or without FGFR inhibitor or MEK inhibitor (n=3). All values are expressed as arithmetic means ± SE.

### 6. cFGF23 mitigates FGF23-mediated inflammatory and immune gene signatures *in vitro*

The presence of Klotho on the surface was crucial for the transcriptomic response to both physiological and pathological FGF23 levels. To investigate whether FGF23 binding to Klotho was specifically involved in this response, we used recombinant human cFGF23 peptide to prevent FGF23 binding to the FGFR-Klotho co-receptor complex^16^ (Fig. 6A). Pretreatment with cFGF23 blocked the FGF23-mediated upregulation of the early-ERK target *EGR1* as well as the late-ERK target *ETV4* in a dose-dependent manner (Fig. 6B). A high dose of cFGF23 (10 μM) alone did not affect *EGR1* mRNA levels; however, it slightly but significantly increased *ETV4* mRNA levels (Fig. 6B, right panel). Next, we validated the inflammation and immune targets such as *TNFRSF9/12A*, *GDF15*, *ANXA1*, and *TGFB1* using qPCR. Similar to ERK targets, FGF23 upregulated these genes, which were partially rescued by cFGF23 co-treatment in a dose-dependent manner (Fig. 6C). Interestingly, cFGF23 alone did not affect the mRNA expressions of any of the inflammatory and immune targets, suggesting that potential “off-target” effects of cFGF23 are probably limited to *ETV4* mRNA (Fig. 6C).

**Fig. 6.**
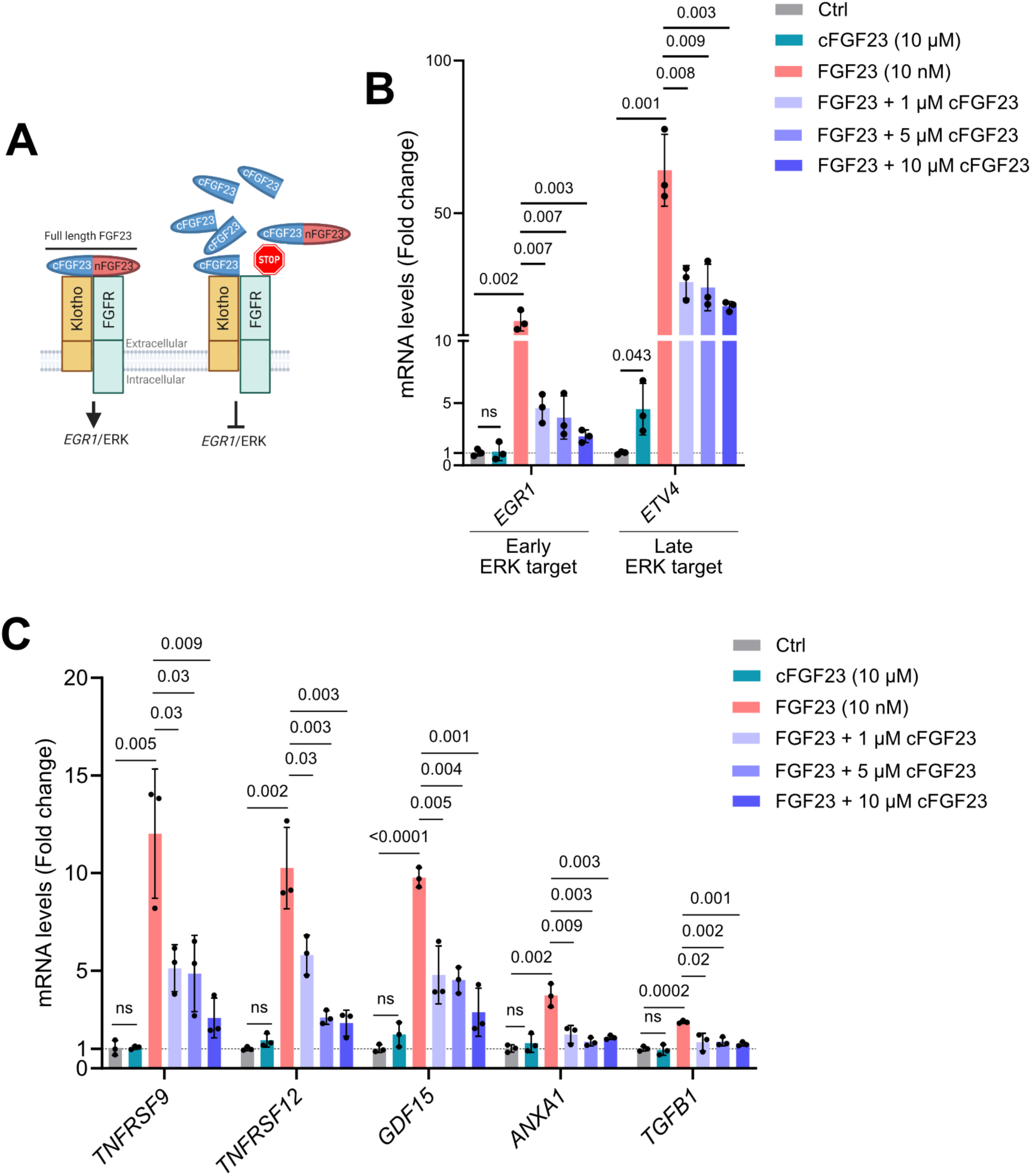
Pathological FGF23-mediated ERK signaling, inflammatory and immune gene signature is mitigated by cFGF23. **A)** Graphical illustration showing mechanisms of action of the cFGF23 peptide. *Left panel:* Intact full-length iFGF23 harbors its C-terminus cFGF23 and N-terminus nFGF23. Within iFGF23, cFGF23 and nFGF23 bind with co-receptors Klotho and FGFR, respectively, and activate ERK signaling. *Right panel:* High concentration of the cFGF23 peptide competes with iFGF23 for binding with Klotho and thus mitigates ERK signaling. **B)** *EGR1* and *ETV4* mRNA, normalized to *GAPDH*, upon treating HEK^Kl^ cells with pathological FGF23 (10 nM FGF23 for 24 hours) with or without increasing concentrations of cFGF23 (n=3). **C)** mRNA levels of *TNFRSF9*, *TNFRSF12A*, *GDF15*, *ANXA1*, and *TGFB1* normalized to *GAPDH*, upon treating HEK^Kl^ cells with pathological FGF23 with or without increasing concentrations of cFGF23 (n=3). All values are expressed as arithmetic means ± SE.

**Fig. 7.**
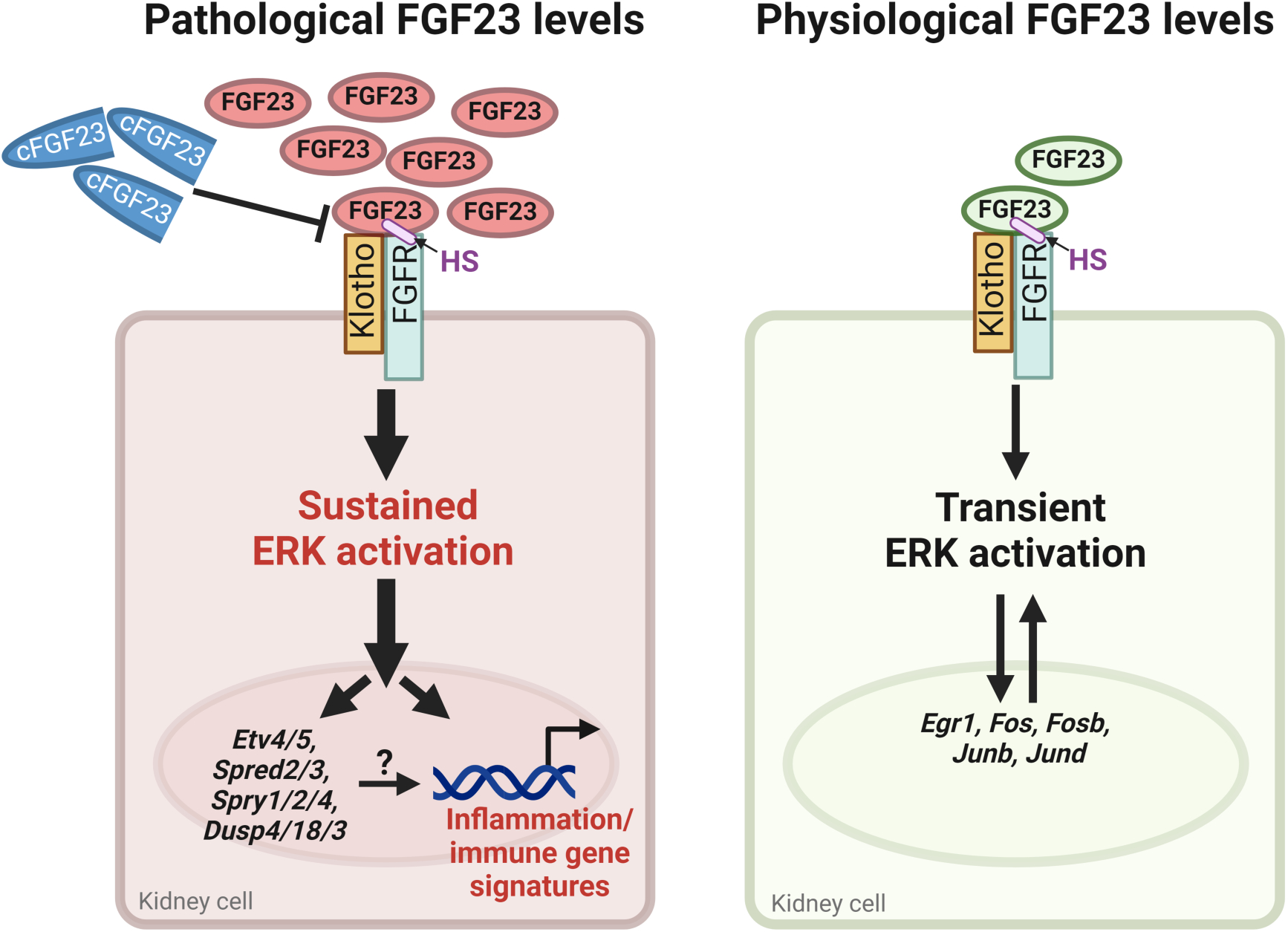
Proposed model: **Left Panel:** In diseases where FGF23 is chronically elevated, the FGF23-FGFR-Klotho-HS signaling complex induces sustained-ERK activation in the cytoplasm. ERK is continuously and irreversibly translocated to the nucleus, where it upregulates late-ERK targets along with inflammation and immune gene signatures. It is currently unclear whether late-ERK targets directly activate inflammation and immune gene signatures. The cFGF23 peptide blocks FGF23-Klotho binding, thereby inhibiting downstream ERK signaling as well as inflammation and immune gene signatures. **Right Panel**: Physiological rise in FGF23 rapidly and reversibly activate ERK in the cytoplasm via FGF23-FGFR-Klotho-HS signaling complex. In this case, ERK transiently translocates to the nucleus and upregulates early-ERK targets and is less likely to regulate gene signatures implicated in inflammation and immune responses.

## DISCUSSION

Our study provides three salient findings: ***(i)*** chronically elevated FGF23 levels in mice induce sustained-ERK activation and inflammation in the kidney; ***(ii)*** cFGF23 gene therapy mitigates sustained-ERK signaling and associated inflammatory responses by blocking FGFR-Klotho binding; and ***(iii)*** FGF23 levels drive temporal ERK signaling dynamics *in-vitro*, with acute physiological levels triggering transient-ERK activation and chronic pathological levels inducing sustained-ERK activation and inflammatory gene signatures.

We found that the early-ERK targets were unchanged in the kidneys of Hyp-Duk mice, while the Hyp-Duk kidneys exhibited elevated late-ERK targets (sustained-ERK signaling) along with activation of inflammatory and immune signaling pathways. The structurally related isoforms, ERK1 and ERK2, were the first MAPKs identified, based on early observations of their rapid and reversible phosphorylation in response to growth factor stimulation^42^. Activated ERK translocates into the nucleus and stimulates transcription factors associated with immediate early genes (*Egr*, *Fos*, and *Jun* families)^43–45^. This type of transient-ERK activation is known to participate in FGF23-mediated mineral homeostasis in the kidney^7,8,46,47^. Conversely, chronically high FGF23 levels in Hyp-Duk mice elevated late-ERK targets (*Etv*, *Spred*, *Spry* families) likely due to continuous and irreversible translocation of ERK into the nucleus. The sustained-ERK signaling is known to trigger inflammatory and immune responses by activating transcription factors like Nuclear Factor kappa-light-chain-enhancer of activated B cells (NF-κB), ETS-like gene 1 (Elk1), cAMP response element-binding protein (CREB), Signal transducers and activators of transcription (STAT)^39,40,48^. Consistently, genes associated with inflammatory and immune cell infiltration, including *Adgre1* (F4/80), *Ptprc* (CD45), and *H2-Aa* (MHC-II), were significantly upregulated in the kidneys of Hyp-Duk mice and may contribute to the proinflammatory responses. Future studies integrating spatial and protein-level analyses will further refine the mechanistic framework proposed here.

Our findings are consistent with recent elegant single-cell transcriptomic analyses demonstrating Klotho-dependent FGF23 bioactivity in discrete nephron segments and its association with inflammatory gene programs^49^. Present study extends these observations by identifying sustained ERK activation as a mechanistic driver of proinflammatory transcriptional responses under elevated FGF23 conditions and demonstrates that these effects can be mitigated through cFGF23 gene therapy.

Renal NGAL expression and its urinary levels, a marker of kidney injury and inflammation, were significantly elevated in Hyp-Duk mice and were rescued by cFGF23 gene therapy. Classical fibrosis markers, such as *Col1a1* and *Col3a1*, were also upregulated, whereas the conventional kidney injury marker *Havcr1* (KIM-1) was not elevated (table S1), suggesting subclinical kidney injury. Consistently, as reported in prior studies, plasma BUN and creatinine levels were unchanged in Hyp mice^50,51^. This implies that elevated FGF23 alone is probably inadequate to cause renal insufficiency without a second hit, such as aging, hypertension, hyperphosphatemia, uremia, or other CKD-related factors. While this manuscript was in preparation, Bockmann and colleagues reported significant renal comorbidities in pediatric XLH patients, including reduced eGFR and elevated urinary markers of glomerular and tubular injury and inflammation^52^. Both XLH patients and mouse models with elevated FGF23 may exhibit subclinical or undetected kidney injury driven by sustained-ERK signaling and associated inflammation.

The convergence of sustained-ERK and inflammatory gene signatures across HEK^Kl^ cells and Hyp-Duk kidneys supports the notion that elevated FGF23-FGFR-Klotho signaling in HEK^Kl^ cells recapitulates, at least in part, the renal transcriptional program observed in Hyp-Duk mice. Despite HEK293 cells being a transformed, non-renal cell line, the transcriptomic response to pathological FGF23 was conserved. This is consistent with recent single-cell transcriptomic study demonstrating that Klotho-dependent FGF23 signaling in HEK^Kl^ cells recapitulates key renal transcriptional programs, including MAPK activation and inflammatory gene induction^49^. Given the absence of renal cell models with native Klotho, HEK^Kl^ offers as a tractable in vitro system to model renal FGF23-FGFR-Klotho transcriptomics, while acknowledging inherent differences between transformed cell lines and native nephron segments. Notably, the newly identified proinflammatory FGF23 signaling targets in HEK^Kl^ cells (*TGFB1, TNFRSF9/12A8, ANXA19,* and *GDF1510*) are also elevated in human CKD patients^53–56^; however, whether their in vivo induction reflects FGF23 signaling or secondary inflammatory effects remains unclear.

MEK inhibitor rescued FGF23-mediated sustained-ERK signaling as well as inflammatory and immune gene signatures. FGFR inhibitor completely abolished ERK signaling and inflammatory gene signatures. Both Klotho and FGFRs are expressed in renal proximal and distal tubules, participating in the canonical effects of FGF23 in mineral metabolism^10,11,57,58^. FGFR1/3/4 are implicated in FGF23-Klotho signaling^6,13^. Notably, only FGFR4 was significantly downregulated, while FGFR1/3 remained unchanged in the kidneys of Hyp-Duk mice (Fig. S8). The specific FGFRs mediating the inflammatory FGF23-ERK signaling is presently unclear. Previous studies have shown that both FGFR and ERK inhibition in Hyp mice corrected mineral metabolism and skeletal phenotype^47,59^. It is tempting to speculate that these inhibitors might also block the inflammatory gene signatures triggered by FGF23-mediated sustained-ERK signaling in the kidneys.

A hallmark of our study is that cFGF23 mitigates FGF23-mediated sustained-ERK signaling, and inflammatory gene signatures both *in vitro* and *in vivo*. In Hyp-Duk mice, AAV-cFGF23 treatment led to high levels of human cFGF23 in mouse serum, effectively correcting inflammatory and immune gene signatures. The structure of the FGF23-FGFR-Klotho complex shows that the C-terminal region of FGF23 binds into an extended groove between the Kl1 and Kl2 domains of Klotho^6,13^. The isolated cFGF23 peptide can bind specifically into this groove, competitively inhibiting Klotho-dependent FGF23 signaling^16^. Thus, cFGF23 likely mitigated Klotho-dependent FGF23 actions in Hyp-Duk mice. Supporting our findings, a previous study showed no change in inflammation-related gene expression in the heart and liver of Hyp mice, where Klotho is not expressed^33^. These findings corroborate our *in vitro* data, concluding Klotho is needed for FGF23 signaling. There was no significant transcriptomic response to FGF23 in HEK^Wt^ cells, and ERK activation and inflammatory/immune gene signatures were observed only in HEK^Kl^ cells. These findings largely align with recent studies demonstrating that Klotho is essential for FGF23 signaling^6,13,49^. On the other hand, elevated FGF23 levels were previously shown to trigger inflammation in extrarenal cells such as immune cells and hepatocytes, independent of Klotho^26,28–30^. At present our findings cannot refute Klotho-independent FGF23 actions in inflammation^20,41^. Moreover, recent studies demonstrating direct biological actions of cFGF23 peptides during inflammation underscore the evolving and context-dependent complexity of FGF23 cleavage biology^60^.

The FGF23-mediated proinflammatory sustained-ERK signaling proposed here may have significant implications in human CKD. Preclinical studies suggest that MEK Inhibitor ameliorates CKD by suppressing ERK signaling^61–63^. It remains to be investigated whether the effect was specifically due to blocking FGF23-mediated sustained-ERK signaling. Our study suggests that the beneficial effects of MEK inhibitor in CKD, at least partly, are mediated by blocking FGF23-mediated sustained-ERK pathway. Importantly, recent studies demonstrated that cFGF23 treatment could improve anemia, heart function and some kidney functions in murine models of CKD^23,64,65^. Gene therapy with AAV-cFGF23 may provide a better efficacy due to its known capacity to release steady levels of cFGF23 in circulation without the peak-and-trough effect known to be associated with protein therapy^17^.

In conclusion, we propose a model (Fig. 6) in which, pathologically elevated FGF23 levels trigger inflammatory and immune gene signatures through the FGFR-Klotho complex and sustained-ERK activation. These untoward effects of FGF23 were significantly mitigated by cFGF23 gene therapy, warranting further investigation to confirm its clinical potential.

## MATERIALS AND METHODS

All the chemicals were purchased from Sigma Aldrich unless otherwise stated.

### Culture of HEK293 cells

HEK293 cell line (HEK^Wt^) was originally purchased from ATCC. The cells were cultured in a growth medium consisting of Minimum Essential Medium MEM, with L-Glutamine (gibco; Cat #11095-080) supplemented with 10% heat-inactivated FBS and 100 U/ml penicillin, and 100 μg/ml streptomycin (gibco; Cat. #15140122). Cells between passages 6 and 17 were used in this study.

### Generation of HEK293^Kl^ cells

For deriving a stable Klotho cell line, HEK^Wt^ cells were transfected with 3 μg of plasmid DNA (Addgene; Cat #17712; http://n2t.net/addgene:17712; RRID:Addgene_17712) in 60 mm culture dish using Jetprime (Polyplus; Cat #55-134) transfection reagent following manufacturer’s protocol. Cells stably expressing Klotho (HEK^Kl^) were selected using 400 μg/ml of G418. Subsequently, stably expressing Klotho (HEK^Kl^) cells were expanded and tested for Klotho protein and mRNA expression by immunoblotting and qPCR respectively. Then HEK^Kl^ aliquots were frozen at -150°C until further use.

### FGF23 treatments in HEK cells

Both HEK^Wt^ and HEK^Kl^ cells undergo exactly similar experimental conditions. A similar number of HEK^Wt^ and HEK^Kl^ cells were seeded in a 6-well plate. After 48 hours, the cells were serum-starved for 16 hours and treated with either PBS or human recombinant FGF23 (Biotechne; Cat #2604-FG/CF) dissolved in PBS. For RNA-seq analysis, following two protocols were applied:

#### Physiological FGF23 treatment

HEK^Wt^ and HEK^Kl^ cells were treated with 0.5 nM FGF23 or control (PBS) for 1 hour.

#### Pathological FGF23 treatment

HEK^Wt^ and HEK^Kl^ cells were treated with 10 nM FGF23 or control (PBS) for 24 hours. After each experiment, cells were washed with ice-cold PBS, and RNA was isolated as mentioned elsewhere.

### Treatment with FGFR/ERK inhibitor and cFGF23 peptide

Forty-eight hours after seeding HEK^Kl^ cells in a 6-well plate, the cells were serum-starved for 16 hours and pretreated with 100 nM FGFR inhibitor PD173074 (Biotechne; Cat #3044/10) or 5 µM MEK inhibitor U0126 (Biotechne; Cat #1144/5) for 1 hour. Then, the treatment was continued for the next 24 hours with 10 nM FGF23 or control (PBS). Isolated cFGF23 peptide^16^ (1, 5, 10 µM) was similarly pretreated for 1 hour and then continued with 10 nM FGF23 for 24 hours.

### Animal experiments

The animal studies were performed according to French and European legislation on animal care and experimentation (2010/63/EU) and were approved by the local Institutional Ethical Board (protocol number 2019-004). In short, the Hyp-Duk mice were obtained from the Jackson Laboratory (Bar Harbor, ME). The treatment consisted of an intravenous injection of AAV-cFGF23 administered at a dose of 1 × 10¹² vg per mouse at 4 weeks of age^17^. The experiment included three groups of mice: PBS-treated WT mice (WT, PBS), PBS-treated Hyp-Duk−/0 mice (Hyp-Duk, PBS), and AAV-cFGF23–treated Hyp-Duk−/0 mice (Hyp-Duk, AAV-cFGF23). Mice were sacrificed after 3 months after intravenous injections of AAV-cFGF23.The cloning of the FGF23 constructs and AAV vector production are described^17^. Urinary NGAL in spot urine samples was measured using a commercial ELISA kit (R&D, Cat. #MLCN20) and normalized to creatinine, measured by the Jaffe method.

To test the species specificity of different FGF23 assays, serum from three male wild-type C57Bl/6 mice (2-3 months old) was spiked with human recombinant FGF23. The FGF23 levels were measured using mouse intact-FGF23 (Quidel, Cat. #60-6800), mouse c-terminal FGF23 (Quidel, Cat. #60-6300), human c-terminal FGF23 (Quidel, Cat. #60-6100) ELISA kits according to the manufacturer’s instructions. We found that only the human cFGF23 ELISA was species-specific and did not detect endogenous mouse FGF23 fragments (Fig. S1B). Both mouse cFGF23 and iFGF23 assays showed cross-reactivity with human FGF23 fragments (Fig. S1C/D).

### Immunoblotting

Both HEK^Wt^ and HEK^Kl^ cells were homogenized in ice-cold RIPA lysis buffer (Thermo Fisher, Cat. #89900) containing protease and phosphate inhibitors. Then cell lysates were centrifuged for 10 min at 2000 g and supernatants were used for immunoblotting. Protein concentration was measured by Bradford assay (CooAssay Protein Dosage Reagent; Uptima, Cat. #UPF86421). Equal amounts of protein (30 μg) were loaded in Laemmli buffer (pH 6.8) on 8% polyacrylamide gels. Electrophoretically separated proteins were blotted to nitrocellulose membranes at 100 V for 2 hours. Next, membranes were blocked in Odyssey blocking buffer (LI-COR; Cat. # 927-70001) for 1 hour. The membrane was further incubated with diluted anti-human Klotho monoclonal antibody/Clone: KM2076 (TransGenic; Cat. #KO603), pERK1/2 (Cell Signaling; Cat #4370S) and tERK1/2 (Cell Signaling; Cat. #4695S) overnight at 4°C. On the following day, diluted secondary antibodies goat-anti-rabbit IRDye 800 (LI-COR; Cat. #926-32211), goat-anti-rat IRDye 800 (LI-COR; Cat. #926-32219) and goat-anti-mouse IRDye 680 (LI-COR; Cat. #926-68070) were added to the membranes in Casein Blocking solution in deionized water (1:10) and incubated for 1 hour at room temperature. Then membranes were repeatedly washed with phosphate-buffered saline containing 0.1% Tween 20. The fluorescent signal was visualized using Odyssey IR imaging system (LI-COR Biosciences).

### RNA isolation and qRT-PCR

The RNA was extracted from HEK^Wt^ and HEK^Kl^ cells using a Nucleospin RNA isolation kit (Macherry Nagel; Cat. #740955). The RNA was transcribed using a High-Capacity cDNA Reverse Transcription Kit (Thermo Fisher; Cat. #4374966) and subjected to qPCR with SybrGreen Master Mix (Roche; Cat. #4707516001) using the following primers:

hKL-Fwd: GCTTTCCTGGATTGACCTTG

hKL-Rev: TGTAACCTCTGTGCCACTCG

hEGR1-Fwd: AGAAGGACAAGAAAGCAGACAAAAGTGT

hEGR1-Rev: GGGGACGGGTAGGAAGAGAG

hETV4-Fwd: TGGAGAGCAGTGCCTTTACTC

hETV4-Rev: TTGATGGCGATTTGTCTGGGG

hTGFB1-Fwd: GCCACAGATCCCCTATTCAA

hTGFB1-Rev: GTCTCAGTATCCCACGGAAA

hTNFRSF9-Fwd: TCTTCCTCACGCTCCGTTTCTC

hTNFRSF9-Rev: TGGAAATCGGCAGCTACAGCCA

hTNFRSF12a-Fwd: CCAAGCTCCTCCAACCACAA

hTNFRSF12a-Rev: TGGGGCCTAGTGTCAAGTCT

hGDF15-Fwd: GAGCTGGGAAGATTCGAACA

hGDF15-Rev: AGAGATACGCAGGTGCAGGT

hGAPDH-Fwd: ACAACTTTGGTATCGTGGAAGGAC

hGAPDH-Rev: CAGGGATGATGTTCTGGAGAG

The relative quantification of gene expression based on double-delta Ct (threshold cycle) analysis was performed after normalization to *GAPDH* expression. All the gene expression values are expressed as arithmetic means ± SD, where n represents a number of independent experiments (biological replicates). An unpaired Student’s t-test was used for comparisons between the groups using GraphPad Prism.

### RNA-sequencing and data analysis

RNA-seq was performed by a commercial vendor (Novogene Company Limited, UK) as previously described^3^. In brief, amplified cDNA samples were subjected to different quality control standards. RNA sample was used for library preparation using NEB Next® Ultra RNA Library Prep Kit for Illumina®. Indices were included to multiplex multiple samples. Briefly, mRNA was purified from total RNA using poly-T oligo-attached magnetic beads. After fragmentation, the first strand cDNA was synthesized using random hexamer primers followed by the second strand cDNA synthesis. The library was ready after end repair, A-tailing, adapter ligation, and size selection. After amplification and purification, the insert size of the library was validated on an Agilent 2100 and quantified using quantitative qPCR. Libraries were then sequenced on Illumina NovaSeq 6000 S4 flow cell with PE150 according to results from library quality control and expected data volume. Differential expression analysis of experimental groups (with different biological replicates per group) was performed using the DESeq2 R package (1.20.0). The resulting p-values were adjusted using the Benjamini and Hochberg’s approach for controlling the false discovery rate. Genes with a padj/p-value<0.05 found by DESeq2 were assigned as differentially expressed. GO enrichment analysis of DEGs was implemented by the clusterProfiler R package. GO terms with padj less than 0.05 were considered significantly enriched by DEGs. Venn diagrams for the sets of genes up- or down-regulated were made by intersecting the lists of gene names between control and FGF23 treatment.

We included 24 tables for *in vitro* RNA-Seq data analysis, three biological replicates in each group. List of eight experimental groups as follows: (1) Kl_FGF23_1h; (2) Kl_Ctrl_1h; (3) WT_FGF23_1h; (4) WT_Ctrl_1h; (5) Kl_FGF23_24h; (6) Kl_Ctrl_24h; (7) WT_FGF23_24h; (8) WT_Ctrl_24h. For *in vivo* RNA-Seq data analysis, five mice in each group were included with three experimental groups as follows: (1) WT; (2) Hyp-Duk; (3) cFGF23-Hyp-Duk. The following thresholds for DEGs were applied: padj<0.05 for *in vitro* data and p-value<0.05 for *in vivo* data. This was due to larger biological variations in mice and the inclusion of five mice per group in the *in vivo* study, compared to three samples per group in the *in vitro* study.

#### Analysis of immune related genes

For the analysis of immune-related genes differentially expressed due to pathological FGF23 treatment *in vitro*, we retrieved a list of curated genes from the InnateDB website (https://www.innatedb.com), which corresponds to a set of genes derived from the Immunology Database and Analysis Portal (ImmPort). Briefly, we intersected the ImmPort gene list with the DEGs from pathological FGF23 treatment in HEK^Kl^ cells (padj<0.05 and abs(log2FC)>1) and used the retrieved gene set for heatmap representation. A similar analysis was performed for the *in vivo* data by translating the ImmPort human genes to their mouse orthologs using biomaRt (version 2.44.4) and intersecting the list with the set of DEGs between Hyp-Duk and WT mice (p-value<0.05). The proportions of immune-related genes rescued by AAV-cFGF23 treatment were obtained by identifying genes overlapping with the ImmPort list that were significantly regulated (p-value<0.05) in opposite directions for the comparisons Hyp-Duk vs. WT and cFGF23-Hyp-Duk vs. Hyp-Duk. The normalized expression values for WT, Hyp-Duk and cFGF23-Hyp-Duk were used for the heatmap.

#### Analysis of biological processes recovered upon AAV-cFGF23 treatment in vivo

For the analysis of GO terms rescued after AAV-cFGF23 treatment, we first retrieved the list of significantly enriched GO terms (padj<0.05) linked to upregulated genes (p-value<0.05) for the comparison of Hyp-Duk vs. WT. The output list was then intersected with the list of enriched GO terms linked to downregulated genes for the cFGF23-Hyp-Duk vs. Hyp-Duk. Representative GO terms included in the intersected output were selected for the barplot representation.

### Deconvolution analysis

Single-cell RNAseq dataset GSE107585 of murine kidney from 7 sex-mixed healthy C57BL/6 mice by Park et al. was downloaded from Gene Expression Omnibus. Using Seurat v 5.3, we excluded cells with less than 200 or more than 2500 features and cells with >5% mitochondrial RNA content. Following normalization and clustering, clusters were annotated based on markers with reference to the original dataset description (Park et al., Science 2018). Bisque v.1.0.5 was then used for deconvolution in reference-based decomposition mode.

### Statistics

All the values are expressed as arithmetic means ± SEM, where n represents number of independent experiments (biological replicates). An unpaired Student’s t test was used for comparisons between the groups using GraphPad Prism. In cases where the p value is not mentioned, the following applies: ns (not significant) p > 0.05, ∗p ≤ 0.05, ∗∗p < 0.01, and ∗∗∗p < 0.001.

### Illustrations

Graphical illustrations were created using a licensed version of Biorender software.

## DATA AVAILABILITY

All the relevant data associated with the current study can be found in the main text or supplementary tables. The raw RNA-seq data used in this publication have been deposited in NCBI Gene Expression Omnibus and are accessible through GEO series accession numbers GSE248734 and GSE269823. All data are available from the authors upon reasonable request.

## ACKNOWLEDGMENTS

This work was supported by the Swiss National Science Foundation through the National Center of Competence in Research NCCR Kidney.CH (N-403-03-55 to G.P.), City University of Hong Kong (9610769 to G.P.), Clinical Research Priority Program HYpertension REsearch NEtwork (CRPP HYRENE) of the University of Zurich (to J. L.), National Natural Science Foundation of China (82073705 & 82273842 to G.C.), Natural Science Foundation for International Senior Scientists (32350710196 to M.M.), Oujiang laboratory startup fund (OJQD2022007 to M.M.), Kungpeng Action Plan award (to M.M.), Natural Science Funding of Zhejiang Province (LR22H300002 to G.C.). The authors thank Prof. Orson Moe and Johanne Pastor from the University of Texas Southwestern Medical Center, Dallas, Texas, USA, for their critical feedback on the manuscript.

## AUTHORS CONTRIBUTIONS

GP conceptualized the study; ASB, CB, GR and GP designed the study; ASB, CB, KK, LJ, FC, AF and GP performed the study; GC, MM and JL contributed new reagents/analytic tools; ASB, CB, KK, LJ, FC, AF GC, MM, CS, LS, JL, GR and GP analyzed the data; GP wrote the manuscript; ASB, CB, KK, LJ, FC, AF GC, MM, CS, LS, JL and GR reviewed and edited the manuscript; GP and JL were responsible for funding acquisition; GP was responsible for supervision.

