## Supplemental information for "Chronically elevated FGF23 drives sustained renal ERK signaling and inflammatory transcriptional programs mitigated by cFGF23 gene therapy"

**Competing Interests**

L.J. and G.R. are inventors in a patent concerning the treatment of XLH (WO/2020/212626). All other authors declare no conflict of interest.

**This pdf contains supplemental material:**

Supplemental figures

Legends for Supplemental tables

### Supplemental Figures

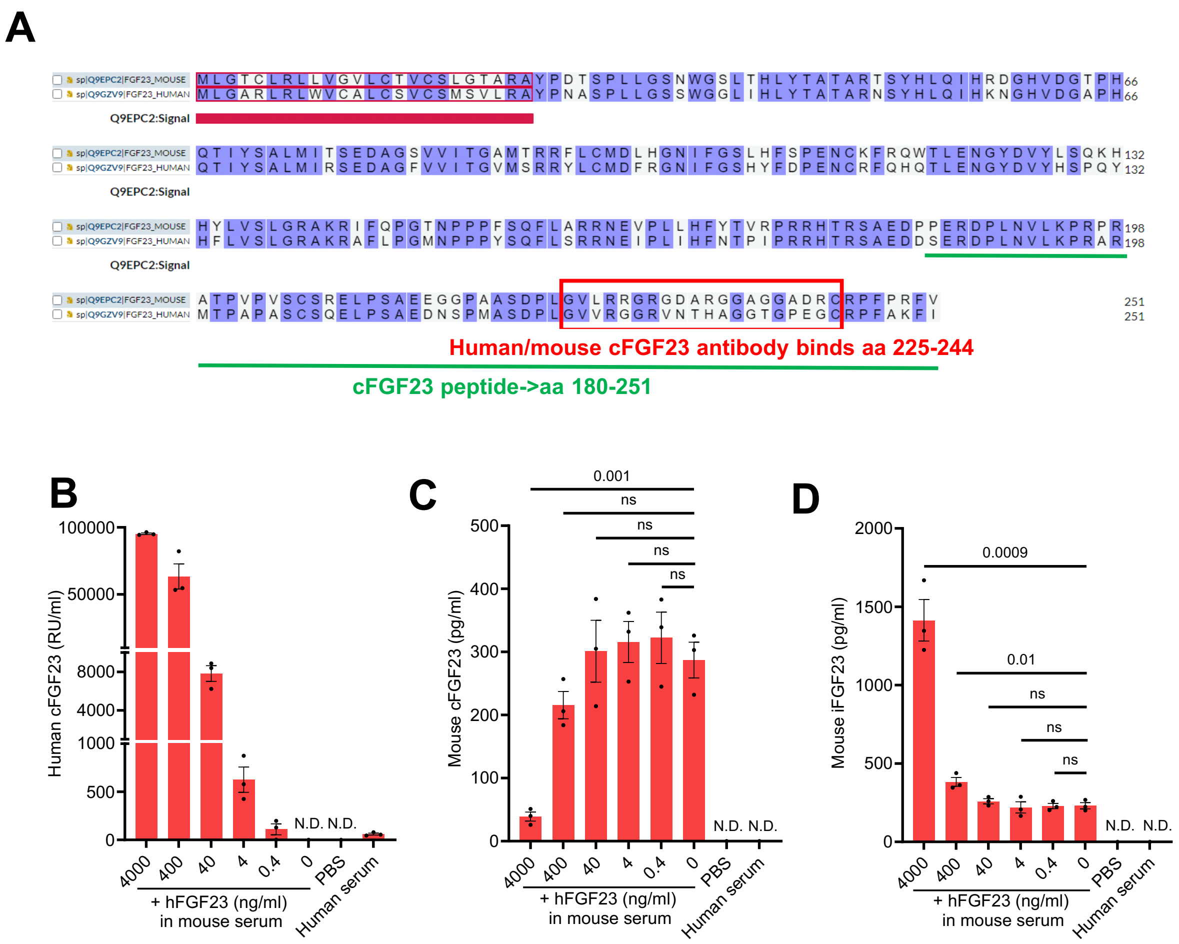


**Fig. S1: Validation of FGF23 assays for species specificity. A)** The amino acid sequences of mouse and human FGF23 (Uniprot) show that most of the sequence is conserved between the two species. However, the region where human and mouse cFGF23 antibodies bind (amino acids 225-244) is relatively less conserved.

Serum from three wildtype C57Bl/6 mice was spiked with human recombinant FGF23 and analyzed using the following ELISA. **B)** The human cFGF23 ELISA specifically detects the human peptide but not the mouse peptide. **C)** The mouse cFGF23 ELISA does not detect the human peptide but shows false negativity when human cFGF23 exceeds 400 ng/ml in mouse serum. **D)** The mouse iFGF23 ELISA does not detect the human peptide at low concentrations but does detect it when human peptide levels exceed 400 ng/ml or more. n=3, each group. ND=not detected. All the values are expressed as arithmetic means ± SE.

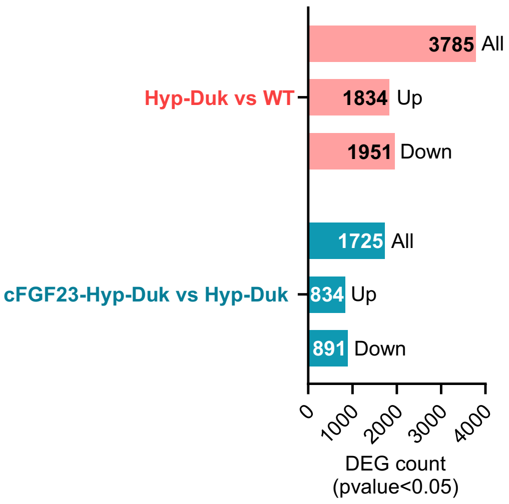


**Fig. S2:** Number and distribution of DEGs in the kidneys of Hyp-Duk versus WT mice and cFGF23-Hyp-Duk versus Hyp-Duk mice.

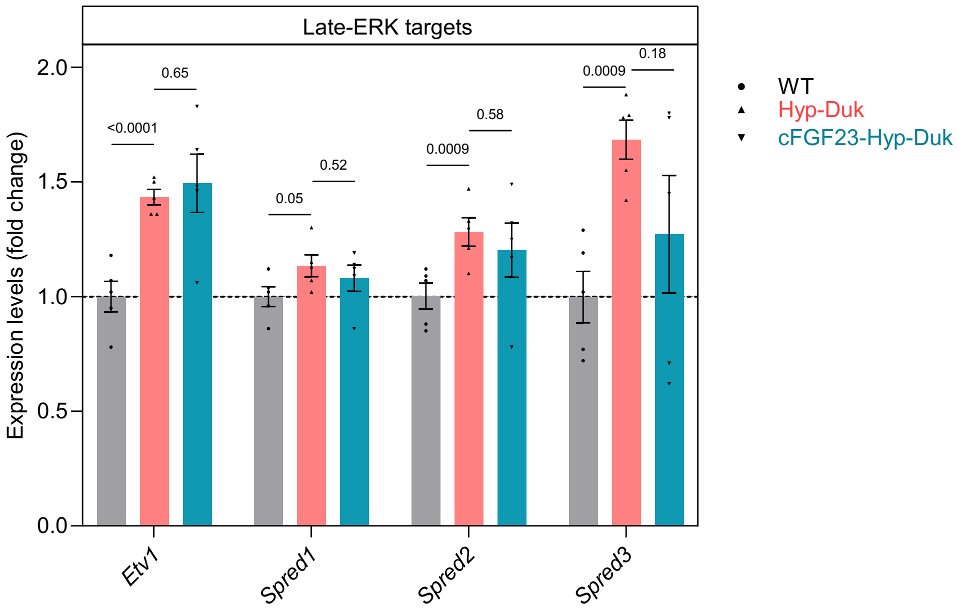


**Fig. S3:** Gene expression levels of sustained-ERK targets in WT, Hyp-Duk, and cFGF23-Hyp-Duk mice. n=5, each group. All the values are expressed as arithmetic means ± SE.

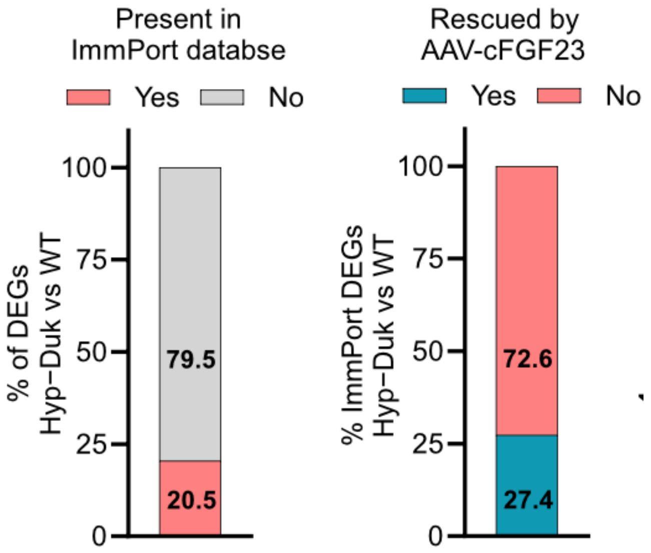


**Fig. S4:** ***Left panel:*** Percentage of DEGs in the kidneys of Hyp-Duk present in the ImmPort database. ***Right panel:*** Percentage of ImmPort DEGs in the kidneys of Hyp-Duk rescued by AAV-cFGF23 treatment.

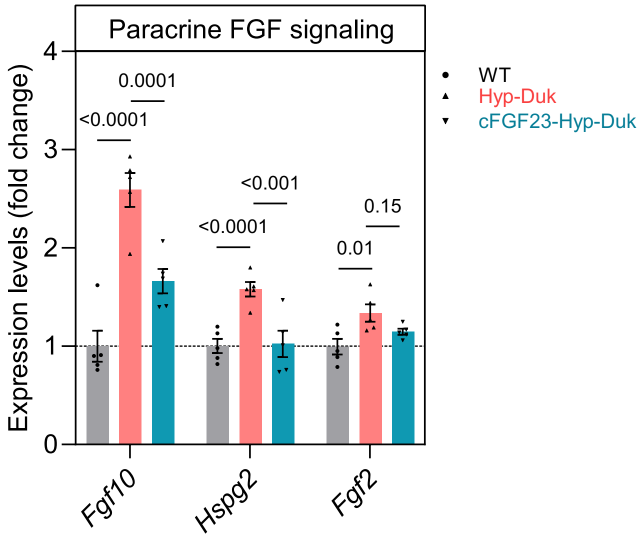


**Fig. S5:** Genes implicated in paracrine FGF signaling in WT, Hyp-Duk, and cFGF23-Hyp-Duk mice. n=5, each group. All values are expressed as arithmetic means ± SE.

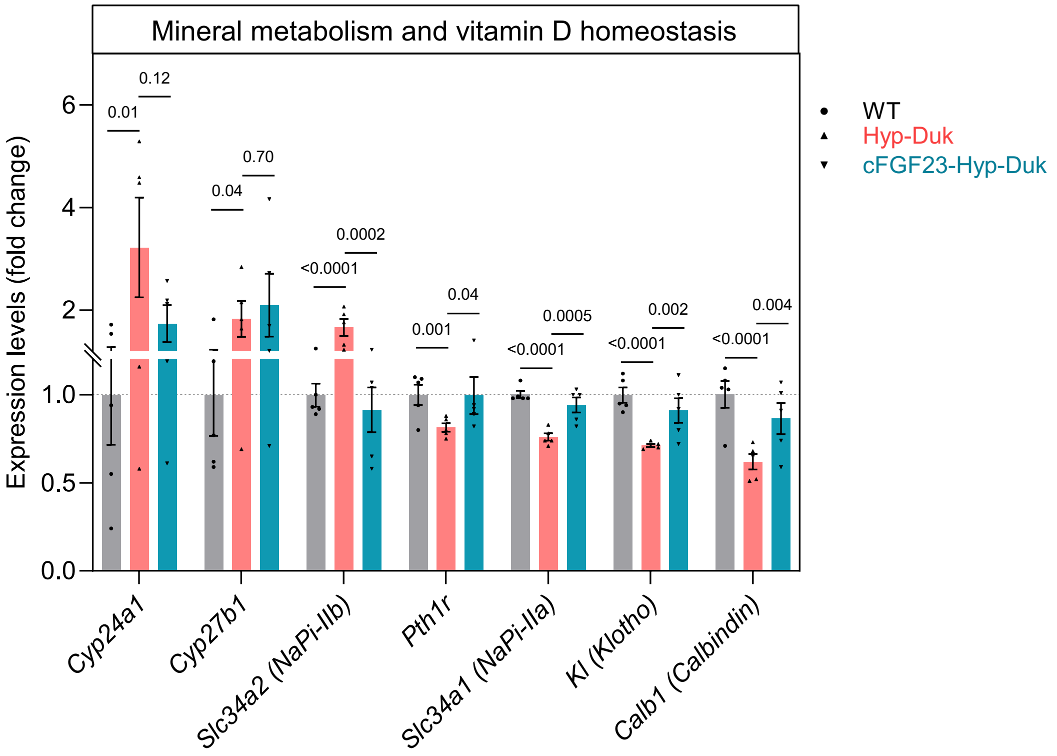


**Fig. S6:** Genes implicated in mineral metabolism and vitamin D homeostasis in WT, Hyp-Duk, and cFGF23-Hyp-Duk mice. n=5, each group. All values are expressed as arithmetic means ± SE.

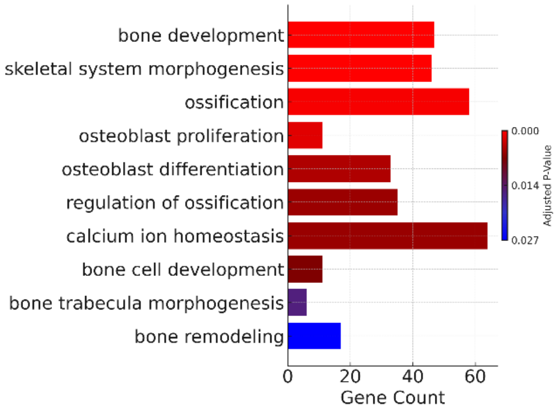


**Fig. S7:** Upregulated GO biological processes altered in kidney samples from Hyp-Duk versus WT mice (padj<0.05) that are associated with skeletal defects.

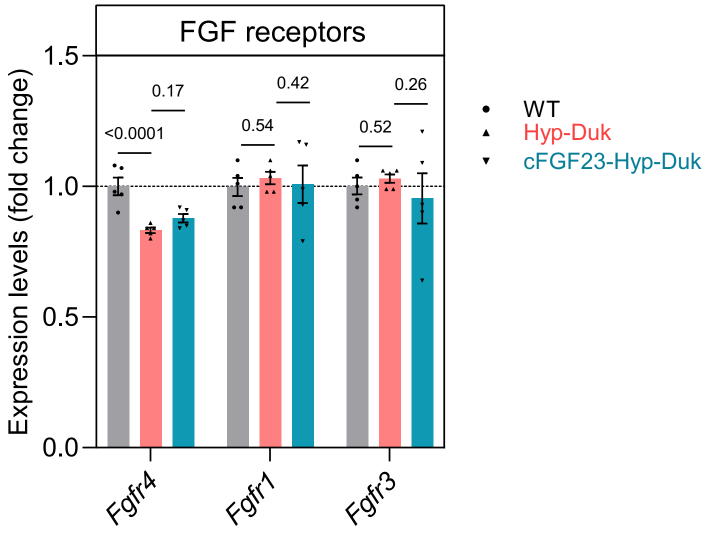


**Fig. S8:** Gene expression levels of various FGFRs in WT, Hyp-Duk, and cFGF23-Hyp-Duk mice. n=5, each group. All the values are expressed as arithmetic means ± SE.

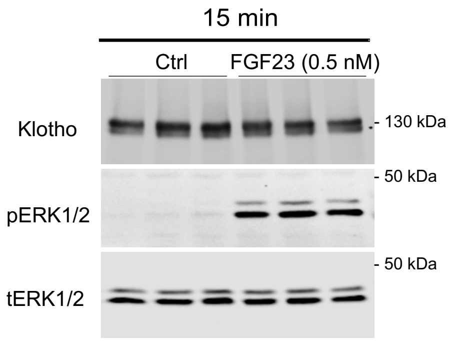


**Fig. S9:** HEK$^{Kl}$ cells respond to FGF23 by upregulating pERK1/2. Original immunoblots show Klotho, pERK1/2 and ERK in HEK$^{Kl}$ cells treated with 0.5 nM FGF23 for 15 minutes.

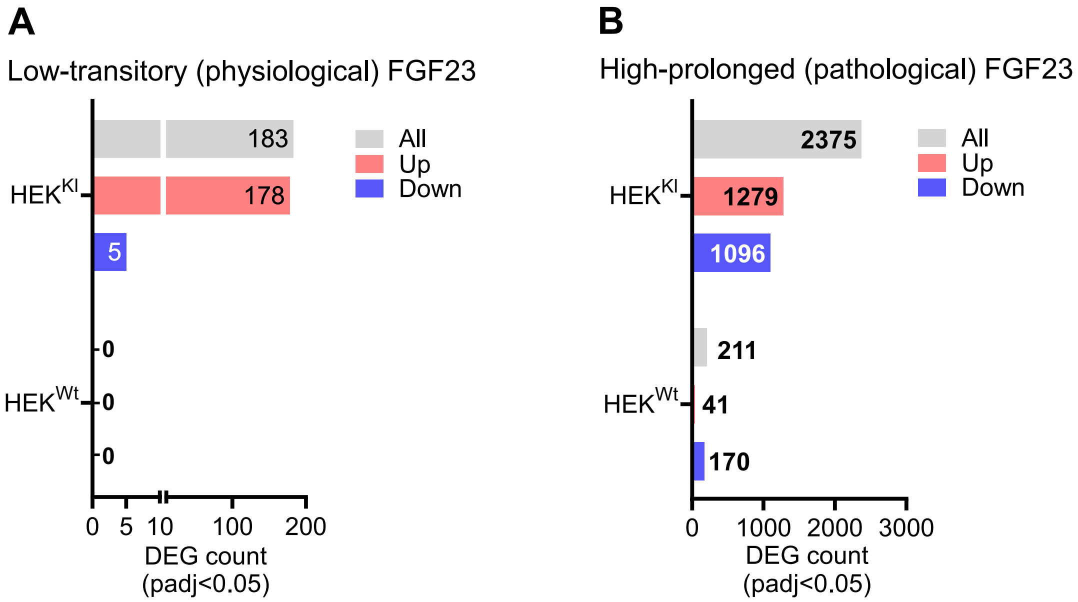


**Fig. S10:** Effect of **A)** low-transitory FGF23 (physiological) and **B)** high-prolonged FGF23 (pathological) on the number of DEGs in $HEK^{KI}$ and $HEK^{Wt}$ cells.

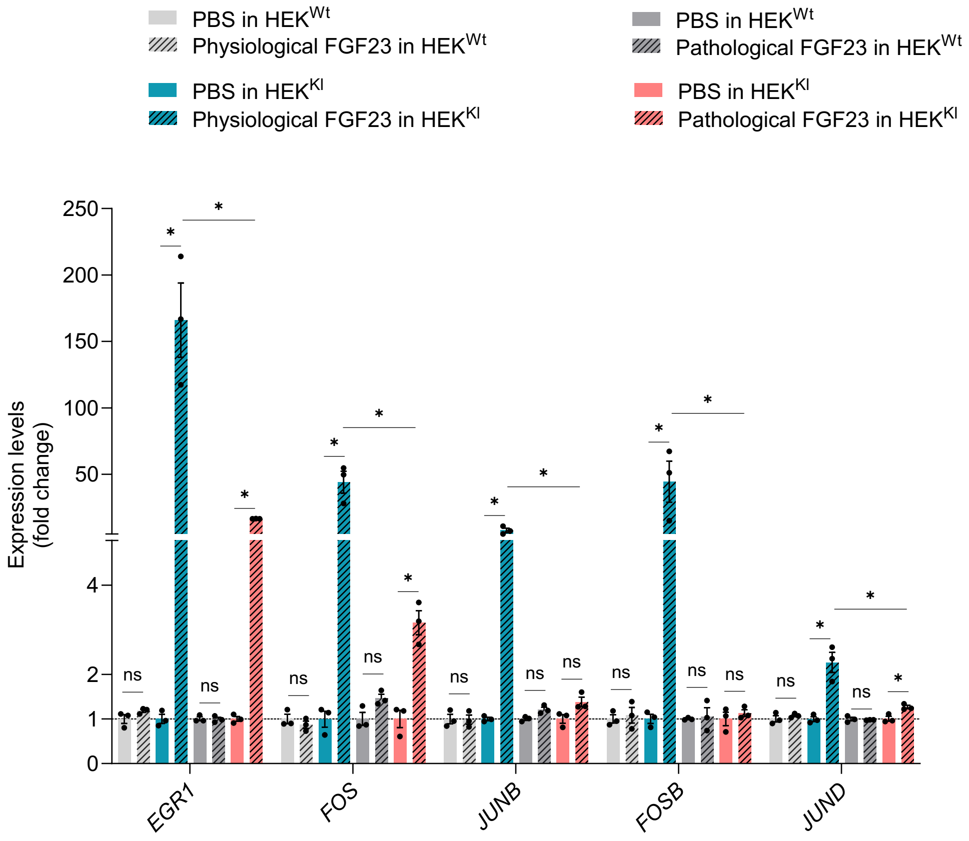


**Fig. S11: Klotho is required for the upregulation of early-ERK targets by physiological FGF23.** Gene expression levels after physiological and pathological FGF23 treatment in $HEK^{Wt}$ and $HEK^{KI}$ cells. n=3, each group. *=padj<0.05, ns=padj>0.05. All the values are expressed as arithmetic means ± SE.

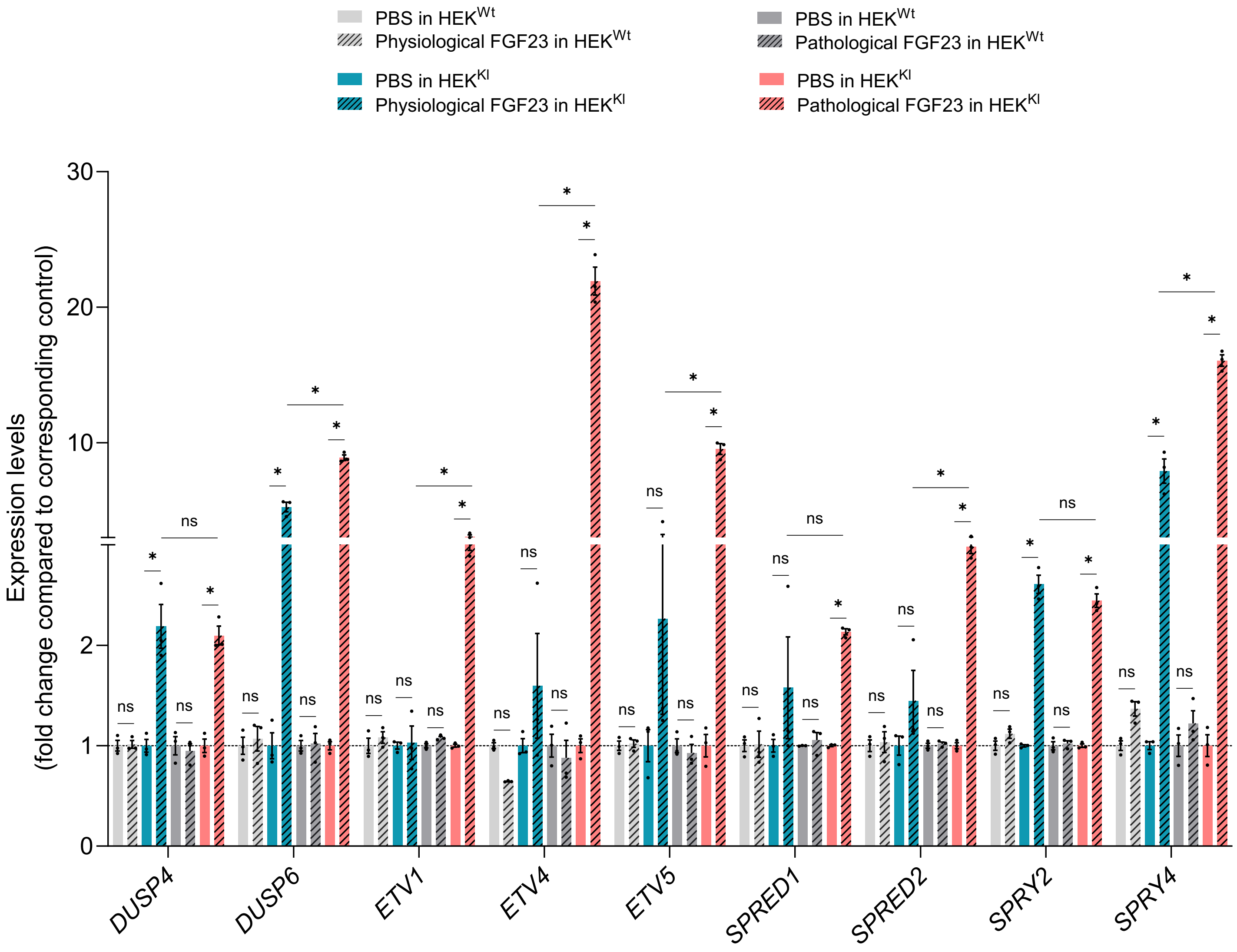


**Fig. S12: Klotho is required for the upregulation of late-ERK targets by pathological FGF23.** Transcripts levels after physiological and pathological FGF23 treatment in HEK$^{Wt}$ and HEK$^{KI}$ cells. n=3, each group. *=padj<0.05, ns=padj>0.05. All the values are expressed as arithmetic means ± SE.

**Legends for supplemental tables:**

**Table S1.**
Differentially expressed genes (DEGs) identified by RNA-seq in kidneys of Hyp-Duk mice compared with wild-type (WT) mice.

**Table S2.**
Differentially expressed genes (DEGs) identified by RNA-seq in kidneys of AAV-cFGF23–treated Hyp-Duk mice compared with untreated Hyp-Duk mice.

**Table S3.**
Gene Ontology (GO) biological processes significantly upregulated in kidneys of Hyp-Duk mice compared with WT mice.

**Table S4.**
Gene Ontology (GO) biological processes significantly downregulated in kidneys of AAV-cFGF23–treated Hyp-Duk mice compared with untreated Hyp-Duk mice.

**Table S5.**
Intersected GO biological processes that are significantly upregulated in Hyp-Duk kidneys and rescued (downregulated) following AAV-cFGF23 treatment, highlighting inflammatory, immune, and ERK-related pathways.

**Table S6.**
Differential gene expression analysis of $HEK^{Wt}$ cells treated with physiological FGF23 (0.5 nM, 1 h) compared with vehicle control.

**Table S7.**
Differential gene expression analysis of $HEK^{KI}$ cells treated with physiological FGF23 (0.5 nM, 1 h) compared with vehicle control, identifying early-ERK target genes (padj < 0.05).

**Table S8.**
Differential gene expression analysis of $HEK^{Wt}$ cells treated with pathological FGF23 (10 nM, 24 h) compared with vehicle control.

**Table S9.**
Differential gene expression analysis of $HEK^{KI}$ cells treated with pathological FGF23 (10 nM, 24 h) compared with vehicle control.

**Table S10.**
Genes upregulated in HEKKI cells treated with pathological FGF23 (10 nM, 24 h) compared with HEKWt cells under identical conditions.

**Table S11.**
Gene Ontology (GO) biological processes enriched among genes upregulated by pathological FGF23 treatment in $HEK^{KI}$ cells.
